# X-Pro: A web-based tool to analyze the loss and gain of local interaction upon a point mutation on protein structures

**DOI:** 10.1101/2025.01.22.634007

**Authors:** Aman Viswakarma, Anurag Neelakanteswara, Saravanamuthu Thiyagarajan

## Abstract

Understanding how point mutations affect protein structures is critical for understanding their impacts on a protein’s function and, eventually their role in causing diseases or biological differences. In recent decades, structural inferences of missense mutations have gained significant attention due to their importance in understanding monogenic disorders. Making point mutations in silico is one of the simple bioinformatics tasks. Existing GUI-based open-source tools like PyMOL, Chimera, Discovery Studio, etc., have simplified this task so that it can be done with a few mouse clicks. However, this continues to be a difficult task for untrained biologists. Further, understanding the loss and gain of interactions caused by such mutations remains even more challenging to them. To address this, we have developed a simple web-browser-based application named *X-Pro* wherein a PDB structure, chain ID, residue number, and the target mutation are taken as inputs, along with mutation information. The tool runs PyMOL and LigPlot tools in the back-end, taking the site of mutation as the ligand and returning graphical images of the structure before and after in-silico point mutation, and a table listing the loss and gain of interactions arising from this mutation. The tool is available for users at the URL http://bts.ibab.ac.in/X-Pro.php.

## Introduction

Proteins play vital roles in biological processes whose constituent amino acids determine their three-dimensional structure, and subsequently their function. Interactions of proteins with other molecules like DNA/RNA/cofactors/substrates/ions/copies of itself, etc., also impact their structures. The preferred conformation of a protein is determined by intra and inter-molecular interactions of a protein with itself or with those in its environment. Mutations in DNA can affect a protein’s sequence, thereby its structure and function. These can be caused by a variety of factors, including errors in DNA replication, exposure to radiation or chemicals, certain infections or disorders, and they can also be inherited from parents^[1]^. These mutations are classified into three types namely, synonymous, missense, and non-sense mutations. Synonymous mutations are harmless whereas non-sense mutations are oftentimes detrimental. Some missense mutations may not affect the protein structure or function as amino acids in the wild type and mutants may have similar biochemical natures and can substitute for each other without causing any loss in their functionality. However, many missense mutations can cause the proteins to misfold or become inactive^[2]^ resulting in a spectrum of outcomes, from negligible differences to fatal phenotypes. Missense mutations in enzymes could change the non-covalent interactions of their respective substrates, making them less efficient or inactive.

Understanding how point mutations affect protein structures is critical for understanding how these mutations might impact protein function and cause or be associated with diseases or biological variations. Several computational tools have been developed to predict the impact of point mutations on protein functions using various approaches, including structure-based and/or sequence-based methods. A few examples include I-Mutant^[3]^, PolyPhen-2^[4]^, and SNPs&GO^[5]^. I-Mutant is a web server and standalone software tool designed to predict the effect of point mutations on protein stability. It employs a Support Vector Machine (SVM) algorithm trained on experimentally determined thermodynamic stability data to predict whether a mutation increases or decreases protein stability. This information is vital for understanding how mutations might affect protein folding and overall structure^[3]^. PolyPhen-2 (Polymorphism Phenotyping v2) is a widely used tool for predicting the functional impact of amino acid substitutions on protein structure and function. It utilizes a combination of sequence-based and structure-based features to classify mutations as one of benign, possibly damaging, or probably damaging. PolyPhen-2 integrates information such as sequence conservation, physicochemical properties of amino acids, and structural annotations to make its predictions. This tool is valuable in prioritizing candidate disease-causing mutations and understanding their potential effects^[4]^. SNPs&GO is another computational tool focused on predicting the functional impact of single nucleotide polymorphisms (SNPs) on protein function. It combines sequence-based features with Gene Ontology annotations to classify mutations into functional categories. By considering evolutionary conservation, protein domain information, and functional annotations, SNPs&GO provides insights into how mutations might affect protein interactions, enzymatic activity, or cellular localization^[5]^. To assess the accuracy of this computational method, the results are compared with easily available and popular tools and servers. Notably, offline and web-based tools such as FoldX and DUET offer computational methodologies to assess the impact of missense mutations. FoldX constitutes an empirical force field designed to swiftly assess the impact of mutations on the stability, folding, and dynamics of proteins and nucleic acids. Its fundamental operation involves computing the free energy of a macromolecule by leveraging its detailed three-dimensional structure^[6]^. DUET represents an integrated computational strategy designed to predict the impact of missense mutations on protein stability. This approach synergizes the capabilities of mCSM and SDM through a consensus prediction framework, which merges the outputs of these individual methods into a refined predictive model^[7]^.

Despite the availability of all such tools, understanding the gain and loss of interactions due to a point mutation remains a non-trivial task and requires expert interpretations. To this regard, we have developed a user-friendly web-based tool, that takes protein structure coordinates and mutation details as input, and return PyMOL based image files, a table of changes in the local interactions due to the induced mutation as returned by the *LigPlot*, and the resultant coordinates for the user’s perusal. The tool is available on the World Wide Web: http://bts.ibab.ac.in/X-Pro.php

## Materials and Method

The details of the mutation to be analysed are captured from the landing page of the tool and processed further.

### In-silico point mutation

The steps involved in analysing potential rotamers of the target residue and selecting the most favoured are described in the PyMOL-Wiki manual pages (https://pymolwiki.org/index.php/Mutagenesis). Python and shell-based scripts have been developed in-house to automate the above process from the command line and are linked through browser-based web pages. The resulting structure, with the mutated residue’s side chain in its selected rotamer conformation, displayed in the *Licorice* representation can be downloaded in the PNG format from the X-Pro tool. The scripts also generate a new PDB file of the protein structure with the desired mutation available for users to download.

### Evaluation of the structural impact of a mutation

The tool *LigPlot*^*[8]*^, is also used in the backend of the X-Pro server. This tool is specifically designed to analyze ligand-protein interactions. Here, the residue at the site of mutation is taken as the ligand. In-house scripts replace the first field ‘ATOM’ to ‘HETATM’, making *LigPlot* recognize the residue at the mutation site as the ligand. The loss and gain of interactions of the residue under consideration with its neighbouring amino acids and solvent molecules are then collated by *LigPlot. LigPlot* also generates 2D schematic diagrams of both wild-type and mutated protein structures using its default parameters. Prior to this calculation, the WT and mutant protein structures are pre-processed using two tools, namely, HBADD and HBPLUS. These tools assign hydrogen atoms and calculate hydrophobic contacts in the input protein structures. (http://www.csb.yale.edu/userguides/graphics/ligplot/manual/index.html). Both wild-type and mutated protein structures are loaded onto the *LigPlot* tool. HBADD initially identifies all heteroatom groups (non-protein atoms, HET group) in the PDB file. Then it subsequently searches for the HET groups in the HET group dictionary, which is available in the PDB. If it is found, the tool generates a definition of the residue type in HBPLUS format. and writes it a file named “hbplus.rc”. This file is then used again by HBPLUS to calculate the hydrogen bonds and hydrophobic contacts. In other words, HBADD prepares the PDB file for HBPLUS by adding the necessary information about HET groups and HBPLUS calculates hydrogen bonds and hydrophobic contacts in the protein. These two tools are used by *LigPlot* GUI automatically, whereas, in the command line mode, they are run manually.

### Web Interface

The pipeline has been integrated into an Apache web server v 2.4.52 hosted on a computer running Ubuntu 22. The interactive user interface of X-Pro has been designed using a combination of front-end web technologies, including XHTML 1.0 for structuring the content, CSS for styling and layout, and JavaScript to enhance interactivity. The tool has been developed using a combination of PHP, MariaDB, and Python. PHP version 8.1.2 has been used to handle server-side logic, MariaDB version 10.6.11 for database management, and Python version 3.10.6 for the backend functionalities.

## Results

A web-based tool that aids in the structural analysis of a point mutation on a protein structure has been developed and hosted on the World Wide Web. The tool can be accessed via the URL: http://bts.ibab.ac.in/X-Pro.php. The landing page and the input sequence of the tool are shown in Supplementary Figure 1

### Data Input

Users can input data via text tabs or a file browser (Figure S1B). First, a PDB ID is input or a local file is uploaded. The analysis begins once this input is received from the user. The available chain IDs and residue numbers are populated from the PDB file (Figure S1C). The input fields are activated sequentially to ensure a complete set of inputs for every run. After the PDB was read, users can select the chain ID, residue number, and the specific target amino acid designated for mutation in the provided text area. Upon obtaining the input data, users are presented with a summary, outlining the input parameters provided (Figure S1D), and in case a user provides an incorrect PDB ID, residue number, or amino acid, an error message will be returned (Figure S1E).

### In-silico point mutation and analysis

The backend of *X-Pro* interfaces with PyMOL and LigPlot to generate the required results. Upon clicking the “analyze” button, rotamer generation and selection happen through PyMOL modules. Subsequently, the input PDB file and that generated from PyMOL are loaded onto the LigPlot tool. which returns i) an image depicting the site of mutation before and after the mutation (Supplementary Figure 2A and 2B), a two-dimensional image output from the *LigPlot+*^*[8]*^ tool (Supplementary Figure 2C and 2D), and iii) the list of loss and gain of interactions (Supplementary Figure 2E and 2F). Users will have the option to download the interaction summary and detailed results. The mutant PDB file is made available for download in case the user wants to avail of other graphics tools for visualization, analysis, or plotting.

### Tool Assessment

The application of the *X-Pro* tool is demonstrated below using the enzyme N-acetylgalactosamine-6-sulfatase (GALNS) as an example. GALNS specifically targets the glycosaminoglycans (GAGs) - keratan sulfate and chondroitin-6-sulfate^[9]^. The presented *X-Pro* output showcases a mutation in the enzyme, featuring a tryptophan substitution at position 78 against Leucine **(Supplementary Figure 2A and S2B)**. Leu78 shows non-covalent interactions with Pro77 and Ddz79, as well as hydrogen bonds with Ser82. Additionally, it engages in hydrophobic interactions with several other amino acids, Asn106, Cit604, Ala107, Pro81, Asn76, Asn289, and Ser80 which are non-ligand residues. Notably, Asn289 established hydrophobic contacts with Pro81, Pro77, and Leu78 **(Supplementary Figure 2C and 2D)**. Ddz79 was particularly significant as the primary active site of the protein^[10]^. Following the mutation, there are notable changes in the interactions. Firstly, the corresponding atoms of Ala107 and Asn106 gained hydrophobic contact with Trp78. A particularly significant change is observed in Asn106, which increases its hydrophobic contacts dramatically from 2 to 25. This extensive increase in hydrophobic contact could increase steric hindrance and potential deformation of the protein structure, impacting its stability and function. Additionally, Ddz79 gained extra non-bonded contact through hydrophobic interaction with Asn289. Furthermore, Tyr108 developed hydrophobic contacts with Pro81, Ser82, and Asn289. There was a reduction in non-bonded contact, decreasing from 3 to 1. This mutation is also analysed with common prediction servers like FoldX (Schymkowitz et al., 2005), Duet (D. E. V. Pires et al., 2014a), mCSM^[11]^, and SDM^[12]^ and all of these predicted this mutation to have a destabilization effect on the protein structure and function.

## Discussion

Missense mutations on proteins affect their interactions with other molecules, and they have the potential to alter protein structure, function, and stability. Mutations are frequently encountered and might even be beneficial in driving evolutionary adaptations and the emergence of desirable traits. Nevertheless, many mutations have the potential to cause disease, resulting in abnormal phenotypes. The currently available structure-based prediction tools primarily report based on a protein’s fold and stability following mutations and do not list gained or lost interactions. Also, most of the popular prediction tools induce the point mutation retaining all common atoms but it is crucial to explore the possible conformations or rotamers that the mutated amino acid can adopt within the altered environment. These alterations directly impact the local interactions at the site of mutation. The specific location and nature of a mutation are key determinants in the context of their influence on a protein’s function, which can range from enhanced interactions to complete abrogation of function. In particular, when mutations cause point changes in protein sequences, they may lead to changes in polarity or hydrophobicity which are crucial significant factors that influence protein structure and function, ultimately affecting the dynamics of how proteins behave and interact within the cell. After introducing mutations into a protein using any GUI, we often need to investigate whether there are gain or loss interactions on account of the mutation, which so far has been left to the trained eye of the researcher rather than having a systematic method repeatable by any user. Addressing this gap, we have developed the tool X-Pro which provides comprehensive insights on changes in the local interactions on a protein, as a result of a mutation considering clash score that is inherently used by PyMOL. Mutating an amino acid using PyMOL involves selecting a suitable rotamer for the mutated residue. A rotamer library is typically used, which contains pre-calculated preferred side chain conformations for various amino acids^[13]^. Each rotamer represents a distinct spatial arrangement of the mutated residue, reflecting the flexibility and adaptability of the protein in response to the mutation. The selected rotamer is adjusted to minimize clashes with other atoms in the protein structure which ensures that the new side chain has the most stable conformation possible. The default output in PyMOL chooses the rotamer having the least clashes with other atoms in the environment. Once this is achieved, the task that remains ahead is identifying the loss and gain of interactions comparing the wild type and the mutant structures. LigPlot is another popular tool that reports interactions when a ligand binds to a receptor. This tool can also compare the changes in interactions when two different ligands binding to the same receptor are analysed simultaneously. X-Pro makes this tool recognize the residue at the site of mutation as the ligand, both in the wild-type and the mutant structure. From this tool, users can visualize graphical representations of the target residues interactions before and after mutation. Additionally, X-Pro allows users to download the mutated PDB files that can be used in any GUI molecular visualization/plotting tool of users’ choice. The significance of the tool is evident in clinical contexts, where understanding the impact of mutations on protein interactions is crucial for predicting the severity of a monogenic disorder. The web-based tool *X-Pro* will be of use to the scientific community aiding them in getting a structural interpretation of genotype-phenotype correlation.

## Supporting Information

X-Pro Landing page and input forms; X-Pro Mutation analysis

## Acknowledgment

This work is supported by ICMR, Government of India, grant No 33/14/2019-TF/Rare/BMS. Institutional grant to IBAB by the Government of Karnataka, India is acknowledged.

## References

[1] H. Nishi, A. Kimura, H. Harada, Y. Koga, K. Adachi, K. Matsuyama, T. Koyanagi, S. Yasunaga, T. Imaizumi, H. Toshima, T. Sasazuki, Circulation 1995, 91, 2911–2915.

[2] M. Petrosino, L. Novak, A. Pasquo, R. Chiaraluce, P. Turina, E. Capriotti, V. Consalvi, Int J Mol Sci 2021, 22, DOI 10.3390/ijms22115416.

[3] E. Capriotti, P. Fariselli, R. Casadio, Nucleic Acids Res 2005, 33, DOI 10.1093/nar/gki375.

[4] I. Adzhubei, D. M. Jordan, S. R. Sunyaev, Curr Protoc Hum Genet 2013, DOI 10.1002/0471142905.hg0720s76.

[5] E. Capriotti, R. Calabrese, P. Fariselli, P. L. Martelli, R. B. Altman, R. Casadio, WS-SNPs&GO: A Web Server for Predicting the Deleterious Effect of Human Protein Variants Using Functional Annotation, 2013.

[6] J. Schymkowitz, J. Borg, F. Stricher, R. Nys, F. Rousseau, L. Serrano, Nucleic Acids Res 2005, 33, W382–W388.

[7] D. E. V. Pires, D. B. Ascher, T. L. Blundell, Nucleic Acids Res 2014, 42, DOI 10.1093/nar/gku411.

[8] R. A. Laskowski, M. B. Swindells, J Chem Inf Model 2011, 51, 2778–2786.

[9] C. P. Morris, X.-H. Guo, S. Apostolou, J. J. Hopwood, H. S. Scott, Genomics 1994, 22, 652–654.

[10] Y. Rivera-Colón, E. K. Schutsky, A. Z. Kita, S. C. Garman, J Mol Biol 2012, 423, 736–751.

[11] D. E. V. Pires, D. B. Ascher, T. L. Blundell, Bioinformatics 2014, 30, 335–342.

[12] A. P. Pandurangan, B. Ochoa-Montaño, D. B. Ascher, T. L. Blundell, Nucleic Acids Res 2017, 45, W229–W235.

[13] L. Dicks, D. J. Wales, J Phys Chem B 2022, 126, 8381–8390.

