## Supporting Information for "X-Pro: A web-based tool to analyze the loss and gain of local interaction upon a point mutation on protein structures"

Dr. Saravanamuthu Thiyagarajan

Associate Professor

Institute of Bioinformatics and Applied Biotechnology (IBAB)

Electronic City Phase 1

Bengaluru – KA 560100

India

**Table of Contents**

| **Page No** | **Item** | **Description** |
| --- | --- | --- |
| 2 | Figure 1 | X-Pro Landing page and input forms |
| 3 | Figure 2 | X-Pro analysis |

B

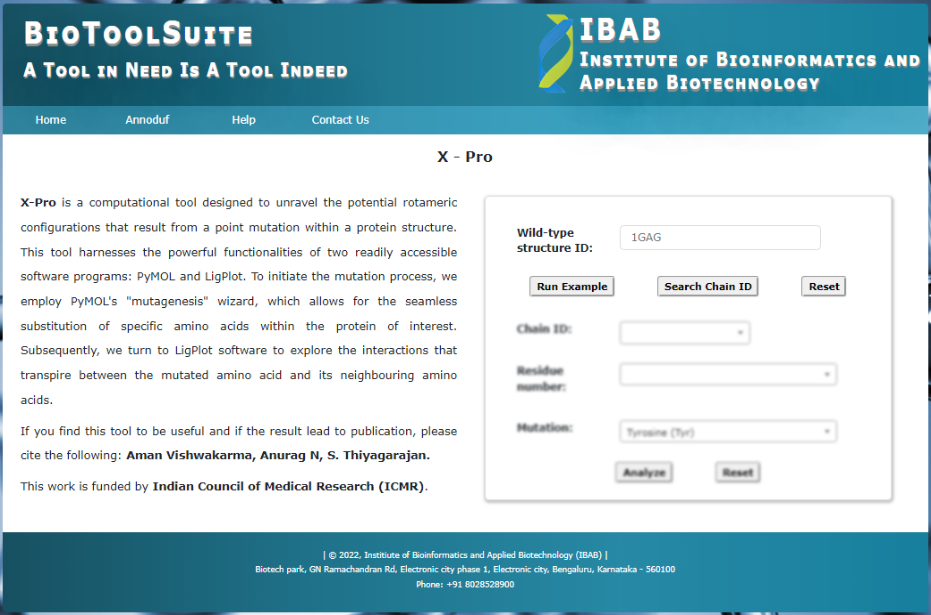

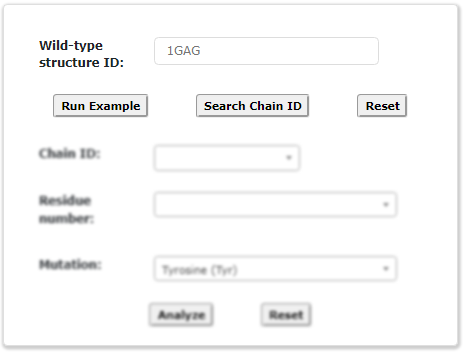

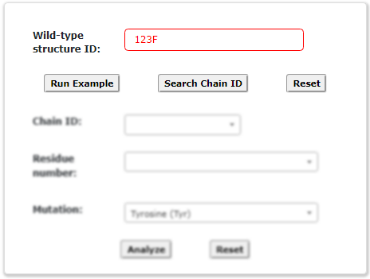

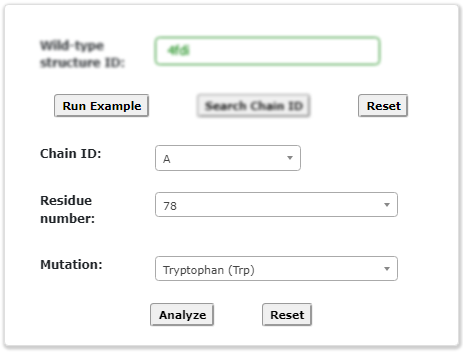

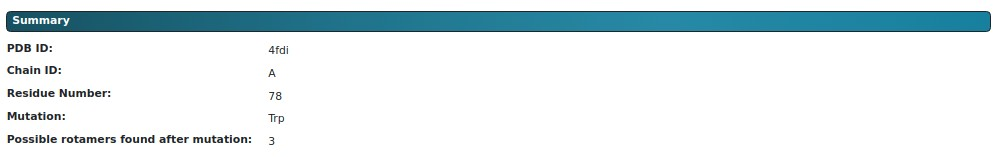

CX

A

D

E

**Figure 1: X-Pro Landing page and input forms:** (A) The screenshot of landing page of the X-Pro web-tool. (B) Coordinates can be input either by giving a PDB id or uploading a local file. Until the file is read and chain IDs get analysed, other fields remain disabled and smeared. (C) upon successful search of PDB, chain ID and residue numbers are populated from the structure. During this, the input structure-ID field is disabled and looks smeared. Upon successful data input, a summary is displayed (D) else the input field turns red (E).

F

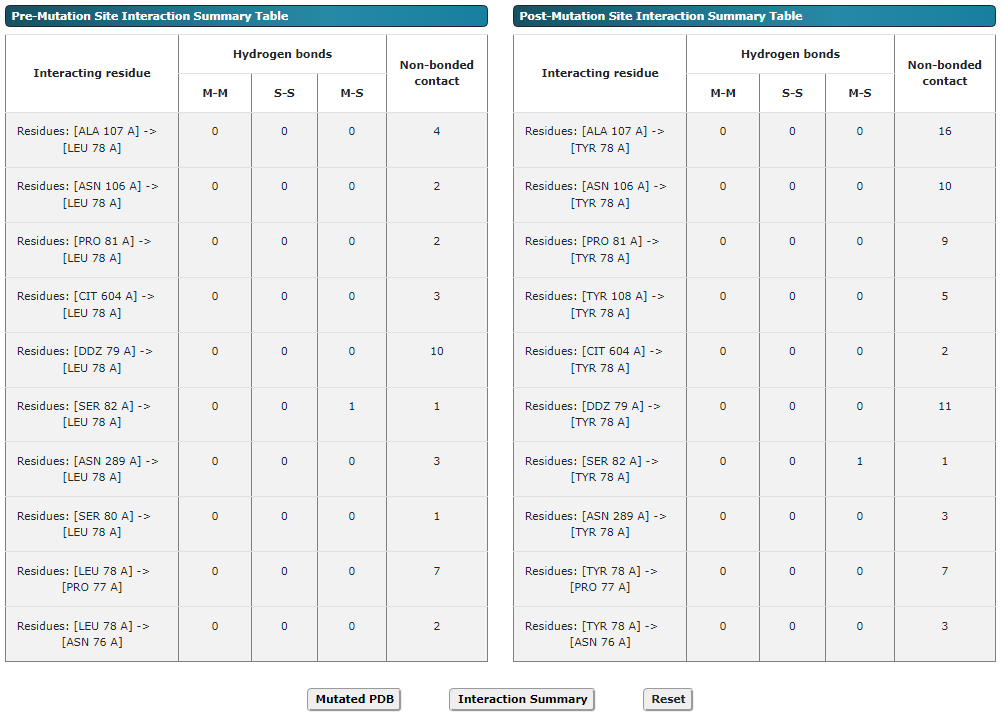

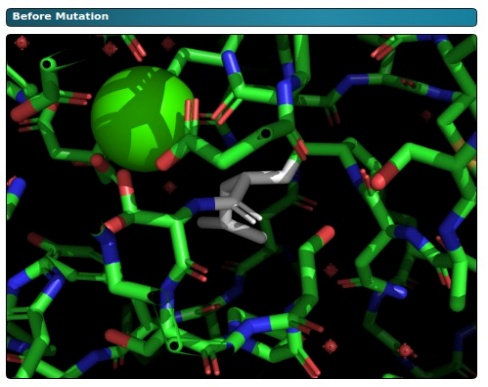

Leu78

Ca

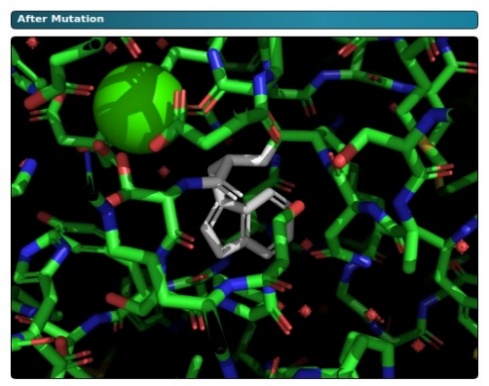

Ca

Trp78

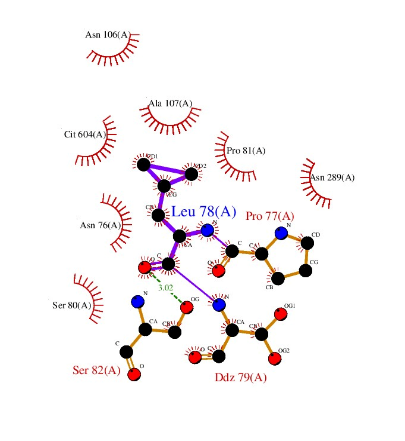

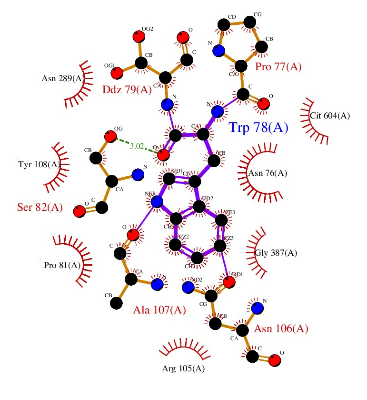

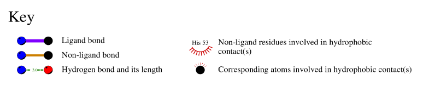

A

B

C

D

E

**Figure 2:X-Pro analysis:** Zoomed-in view of stick representation at the site of mutation, the residue to be mutated is show in grey sticks.
